# TIMS^2^Rescore: A DDA-PASEF optimized data-driven rescoring pipeline based on MS^2^Rescore

**DOI:** 10.1101/2024.05.29.596400

**Authors:** Arthur Declercq, Robbe Devreese, Jonas Scheid, Caroline Jachmann, Tim Van Den Bossche, Annica Preikschat, David Gomez-Zepeda, Jeewan Babu Rijal, Aurélie Hirschler, Jonathan R Krieger, Tharan Srikumar, George Rosenberger, Dennis Trede, Christine Carapito, Stefan Tenzer, Juliane S Walz, Sven Degroeve, Robbin Bouwmeester, Lennart Martens, Ralf Gabriels

## Abstract

The high throughput analysis of proteins with mass spectrometry (MS) is highly valuable for understanding human biology, discovering disease biomarkers, identifying therapeutic targets, and exploring pathogen interactions. To achieve these goals, specialized proteomics subfields – such as plasma proteomics, immunopeptidomics, and metaproteomics – must tackle specific analytical challenges, such as an increased identification ambiguity compared to routine proteomics experiments. Technical advancements in MS instrumentation can counter these issues by acquiring more discerning information at higher sensitivity levels, as is exemplified by the incorporation of ion mobility and parallel accumulation - serial fragmentation (PASEF) technologies in timsTOF instruments. In addition, AI-based bioinformatics solutions can help overcome ambiguity issues by integrating more data into the identification workflow. Here, we introduce TIMS^2^Rescore, a data-driven rescoring workflow optimized for DDA-PASEF data from timsTOF instruments. This platform includes new timsTOF MS^2^PIP spectrum prediction models and IM2Deep, a new deep learning-based peptide ion mobility predictor. Furthermore, to fully streamline data throughput, TIMS^2^Rescore directly accepts Bruker raw mass spectrometry data, and search results from ProteoScape and many other search engines, including MS Amanda and PEAKS. We showcase TIMS^2^Rescore performance on plasma proteomics, immunopeptidomics (HLA class I and II), and metaproteomics data sets. TIMS^2^Rescore is open-source and freely available at https://github.com/compomics/tims2rescore.

## Introduction

Proteomics is an invaluable tool for understanding human biology, facilitating the discovery of new disease biomarkers, identifying potential therapeutic targets, and exploring interactions with pathogens or microorganisms (1–4). Various proteomics subfields have emerged that address specific challenges. For example, plasma proteomics tackles the vast dynamic range of protein abundances, immunopeptidomics deals with the non-tryptic nature of immunopeptides combined with varying lengths, i.e. class I 8-12 amino acids and class II 12-25 amino acids, and metaproteomics must address the complexity of multiple species that have highly similar tryptic peptides within each sample (4–6). These challenges all contribute to higher identification ambiguity, stemming from a higher complexity in the acquired data, a larger and more diverse peptide search space, high sequence similarities, or all of the above (7). To overcome these issues, highly sensitive yet specific peptide spectrum identification strategies are required.

Artificial intelligence (AI) has undeniably transformed many research fields, including computational proteomics. AI allows us to predict analyte behavior for almost every step in the liquid chromatography - ion mobility - tandem mass spectrometry (LC-IM-MS/MS) pipeline (8, 9), from peptide retention times to the intensities of peptide fragmentation spectra. Indeed, machine learning tools such as DeepLC (10) and MS^2^PIP (11) can predict these values very accurately and precisely. Recently, such predictors have been shown to be very powerful means to further increase identification performance through data-driven rescoring (12–14). Here, the predictions for each candidate peptide-spectrum match (PSM) are first compared to the observations, and these comparison values are then provided to a semi-supervised machine learning algorithm to rescore the PSMs based on all available information. Data-driven rescoring algorithms such as MS^2^Rescore (12, 15) have been shown to substantially increase identification sensitivity and specificity while maintaining a proper statistical false discovery rate (FDR) control. For example, for immunopeptidomics data, increases in identification rate of over 35% at 1% FDR have been reported (16–18).

Technical advances to LC-IM-MS/MS instruments, exemplified by timsTOF instruments, have also greatly improved identification performance in challenging proteomics subfields. While standard LC-MS/MS systems rely solely on the LC setup and the quadrupole for peptide separation preceding fragmentation, timsTOF instruments incorporate IM for additional ion separation in the gas phase based on collisional cross sections. Moreover, due to the parallel accumulation and serial fragmentation (PASEF) technology, precursor ions are accumulated in the TIMS tunnel before being released sequentially, leading to much improved sensitivity (19). This higher sensitivity is particularly beneficial for detecting low-abundance ions, common in immunopeptidomics or plasma proteomics. Indeed, timsTOF instruments have been shown to substantially boost identification rates for both class I and class II immunopeptides (6, 20, 21), and to allow for much broader plasma protein profiling (22).

We here present TIMS^2^Rescore, a new version of our data-driven rescoring platform MS^2^Rescore, optimized for data dependent acquisition PASEF (DDA-PASEF) data from timsTOF instruments. First, we have trained new timsTOF-compatible MS^2^PIP spectrum prediction models, which were subsequently validated on plasma proteomics, immunopeptidomics, and metaproteomics data sets. Second, to optimally leverage the additional information provided by the IM dimension, we have developed IM2Deep, a deep learning-based peptide collisional cross section (CCS) predictor that uses a similar architecture to our state-of-the-art retention time predictor, DeepLC. As a result, IM2Deep is able to accurately predict CCS values for both unmodified as well as modified peptides, even if those modifications were not seen during training. Third, for an optimal software integration, TIMS^2^Rescore directly accepts Bruker raw mass spectrometry data and search results from Bruker ProteoScape and many other search engines, including MS Amanda and PEAKS. Finally, we evaluated the full TIMS^2^Rescore workflow, including MS^2^PIP, IM2Deep, and DeepLC, on data sets from plasma proteomics, immunopeptidomics and metaproteomics experiments. In all three cases, TIMS^2^Rescore shows substantial increases in identification performance. Thus, TIMS^2^Rescore will enable researchers to obtain a broader and more confident peptide and protein identification coverage for a large variety of applications.

## Methods

### Specialized MS^2^PIP models for timsTOF fragmentation

A new model (timsTOF 2024) was trained following the procedures described in the 2023 MS^2^PIP publication (20). The trypsin, elastase, and class I immunopeptide data that was used to train the original timsTOF 2023 model (PXD046535, and PXD040385) was now supplemented with class II immunopeptides retrieved from Hoenisch Gravel *et al*. (21) (PXD038782). The 505,289 highest scoring peptidoforms across all data sets were retained – considering precursor charge as part of the peptidoform. These were then further separated into a training set (480,024 peptidoforms) and a test set (25,265 peptidoforms) using a stratified division based on the data set to ensure sufficient peptides from each peptide type, i.e. class I, class II, trypsin-digested, and elastase-digested peptides in each subset. All processed data is available on Zenodo at 10.5281/zenodo.11277943 The training set was used to train XGBoost (v1.7.2) models for singly charged b-and y-ions. The Hyperopt (v0.2.7) package was used for hyperparameter optimization, employing a five-fold cross-validation. The full hyperparameter optimizations were logged with Weights and Biases. For model evaluation, not only the highest scoring test peptidoforms, but all PSMs (193,400) for the test peptidoforms (25,265) were retrieved from the search data. For each PSM, the observed b-and y-ion intensities were retrieved, and predictions were made with the 2021 Orbitrap HCD (higher energy collisional dissociation) model, the timsTOF 2023 model, and the newly trained timsTOF 2024 model. Furthermore, the observed intensities for PSMs coming from the same peptidoforms were also correlated to measure the inherent variation in experimental data, providing an estimate of the best possible accuracy that can be achieved with prediction models.

### IM2Deep ion mobility prediction

The deep learning architecture of IM2Deep mirrors that of DeepLC and is described in detail in Bouwmeester *et al*. (10). Briefly, IM2Deep employs a convolutional architecture with four distinct paths, through which each encoded peptide is propagated. Three paths utilize convolutional and maximum pooling layers to capture local structures. These paths handle atomic composition of amino acids, atomic composition of diamino acids and one-hot encoding for unmodified amino acids. A fourth path passes on global features through densely connected layers, including length, total atomic composition, and composition at specific positions.

The sole difference between IM2Deep and DeepLC is the addition of five features to the global feature matrix: 1) the relative frequency of histidine within the peptide sequence, 2) the relative frequency of bulky amino acids (F, W, Y), 3) the relative frequency of acidic amino acids (D, E), 4) the relative frequency of lysine and arginine (K, R), and 5) the charge state of the peptide ion. These features were included because, as has been shown earlier, groups of amino acids with similar physico-chemical properties can have a similar impact on the CCS and are thus grouped (23). The model combines the results from all paths through flattening and concatenation, providing an input for six connected dense layers in the final combined path, which outputs the predicted CCS value.

To train and evaluate IM2Deep on its ability to generalize its predictions on modifications and amino acids unseen during training, two data sets were combined into one large data set. The first data set, described by Meier *et al*. (23) consists of 718,917 unique combinations of peptide sequence, charge state and, when applicable, modifications (limited to methionine oxidation, cysteine carbamidomethylation and N-terminal acetylation). The second data set, described in (24), comprises 5,202 unique peptidoform-charge state combinations, and contains a wider variety of modifications. In this data set, a distinction is also made between symmetrical and asymmetrical arginine dimethylation. However, as IM2Deep is not able to distinguish between isomeric differences in peptides, we used the mean CCS value of these isomers as the CCS value for the dimethylated peptide-charge state pair. To account for experimental drifts in the measurements of CCS values between the two data sets, we performed an alignment by calculating the linear offset (*y = ax + b*) between overlapping peptide-charge pair states in the two data sets, according to (23) and (25). Only unique peptidoform-charge states were retained in the data set, and the mean value of overlapping pairs, after alignment, was used to train and evaluate the models. Trained models were initialized with random weights drawn from a normal distribution (*μ*=0.0 and *σ*=1.0). A single NVIDIA Geforce RTX 4090 graphic card was used for training, which lasted for maximally 300 epochs, with early stopping on a validation set to prevent overfitting.

To allow the ion mobility dimension to be used for rescoring TIMS data, IM2Deep was implemented as a feature generator within TIMS^2^Rescore. The final IM2Deep model shipped with TIMS^2^Rescore was trained and evaluated (89.1% training, 0.9% validation, 10% test, with no overlap in peptidoforms) on the data set described above, in combination with an immunopeptidomics data set (21) which consists of 437.479 unique (modified) peptide-charge pairs, most of which are non-tryptic. Before merging, the immunopeptidomics data set was aligned to the original data set using a linear offset between overlapping precursors.

The rescoring features generated by IM2Deep include the observed and predicted CCS values, alongside the absolute and percentual error between the observed and predicted CCS. Before computing these features, the predictions are calibrated to the observed CCS range by calculating the linear offset between the CCS values of a reference data set and the overlapping precursors in the 75% most confidently identified precursors at 1% FDR before rescoring.

### Data-driven rescoring

The full TIMS^2^Rescore pipeline, with the new MS^2^PIP and IM2Deep models, was evaluated on unseen plasma proteomics, immunopeptidomics, and metaproteomics data sets. The plasma proteomics data were provided by Bruker. The class II immunopeptidomics runs were generated in-house as well (see Supplementary Methods). The immunopeptidomics class I data was obtained from the same large-scale immunopeptidomics study from Gravnel *et al*. (21) (PXD038782), where the class II data was used for MS^2^PIP training. Lastly, all metaproteomics samples were acquired from the CAMPI study (4) (PXD023217). All data was searched using Sage (v0.14.3>) with 10 ppm precursor and fragment tolerances, maximum 2 variable modifications where in all 4 searches oxidation was considered. For the immunopeptidomics this was further supplied with carbamidomethyl, while this was set as fixed modification for the plasma and metaproteomics. For the immunopeptidomics searches no cleavage rule was used with lengths fixed at 8-25 for class I and 8-30 for class II. The plasma and metaproteomics data set were searched with trypsin as cleavage rule with a restriction for proline, lengths were fixed at 8-50 for plasma and 8-30 for metaproteomics. Aside from the metaproteomics data where the custom sequence database was used that was published alongside the MS data(4), all data sets were searched using the Swiss-Prot canonical human proteome (UP000005640, 20,597 entries, downloaded March 2024).

## Results

### TIMS^2^Rescore: Data-driven rescoring tailored to timsTOF instruments

TIMS^2^Rescore is built on top of the data-driven rescoring framework MS^2^Rescore. Several improvements were made to create a streamlined rescoring workflow that is fully tailored to DDA-PASEF data from timsTOF instruments. First, to drastically speed up reading of large spectrum files, we integrated the Rust-based mzdata file readers for the MGF and mzML file formats (https://github.com/mobiusklein/mzdata). Moreover, we have implemented direct support for Bruker TDF and mini-TDF raw formats, using the TimsRust package (https://github.com/mannlabs/timsrust), allowing users to avoid long data conversion steps altogether. Second, as support for DDA-PASEF data was recently added to the ultra-fast search engine Sage (26), we added direct support for Sage PSM files in TIMS^2^Rescore, along with support for PSM files from the Bruker ProteoScape search environment. Third, a new set of default parameters optimized for rescoring DDA-PASEF data are made available. Together with the new prediction models outlined below, these new features significantly improve the ease of use and computational performance for rescoring timsTOF data.

### MS^2^PIP prediction models

We, and others, have previously shown that different fragmentation methods can heavily alter peptide MS2 spectra (27, 28). We therefore trained new MS^2^PIP models that can accurately predict peak intensities for timsTOF acquired peptides. In 2023, we trained a new model on tryptic peptides, elastase digested peptides, and class I immunopeptides, which we subsequently used to boost coverage of immunopeptides through rescoring (20). While this model performed well for data sets similar to the aforementioned training peptides (median Pearson correlation coefficient (PCC) 0.89, 0.89, 0.87 for tryptic, class I immunopeptides and elastase peptides, respectively), the performance for class II immunopeptides was significantly less (median PCC 0.64) and even outperformed by the model for Orbitrap HCD spectra (Figure 1). This could be due to the generally longer peptides which were not yet seen in training data, as class I immunopeptides and elastase peptides are generally shorter. To overcome this problem, we have trained a new MS^2^PIP model on data that was supplemented with a large amount of class II immunopeptides. The newly trained model performs comparable to the previous timsTOF model for tryptic (0.89 median PCC) and class I immunopeptides (0.88 median PCC), performs slightly worse for elastase digested peptides (0.81 median PCC) but performs drastically better for the class II immunopeptides (0.85 median PCC) (Figure 1). While the performance drops slightly for elastase digested peptides, the predictions accuracy is still close to the expected intensity variance seen in timsTOF spectra, as is also the case for all other peptide types. This drop will likely be due to the lower amount of training peptidoforms in the train set (35,126). However, we chose to retain this peptide type in the trainset to ensure more broadly applicable models. When examining predictions for peptides with a higher variance, we can see that the predictions still approximate the median intensity across all peptide spectra, highlighting the robustness of the newly trained model (Supplementary Figure S1). The drop in performance for the elastase digested peptides could potentially be attributed to the lower amount of training peptides for elastase relative to the newly added class II immunopeptides (35,126 vs 232,798 peptides, respectively). Overall, the 2024 timsTOF model performs similar to, or better than, the timsTOF 2023 model. All median PCCs are listed in Supplementary Table S1.

**Figure 1:**
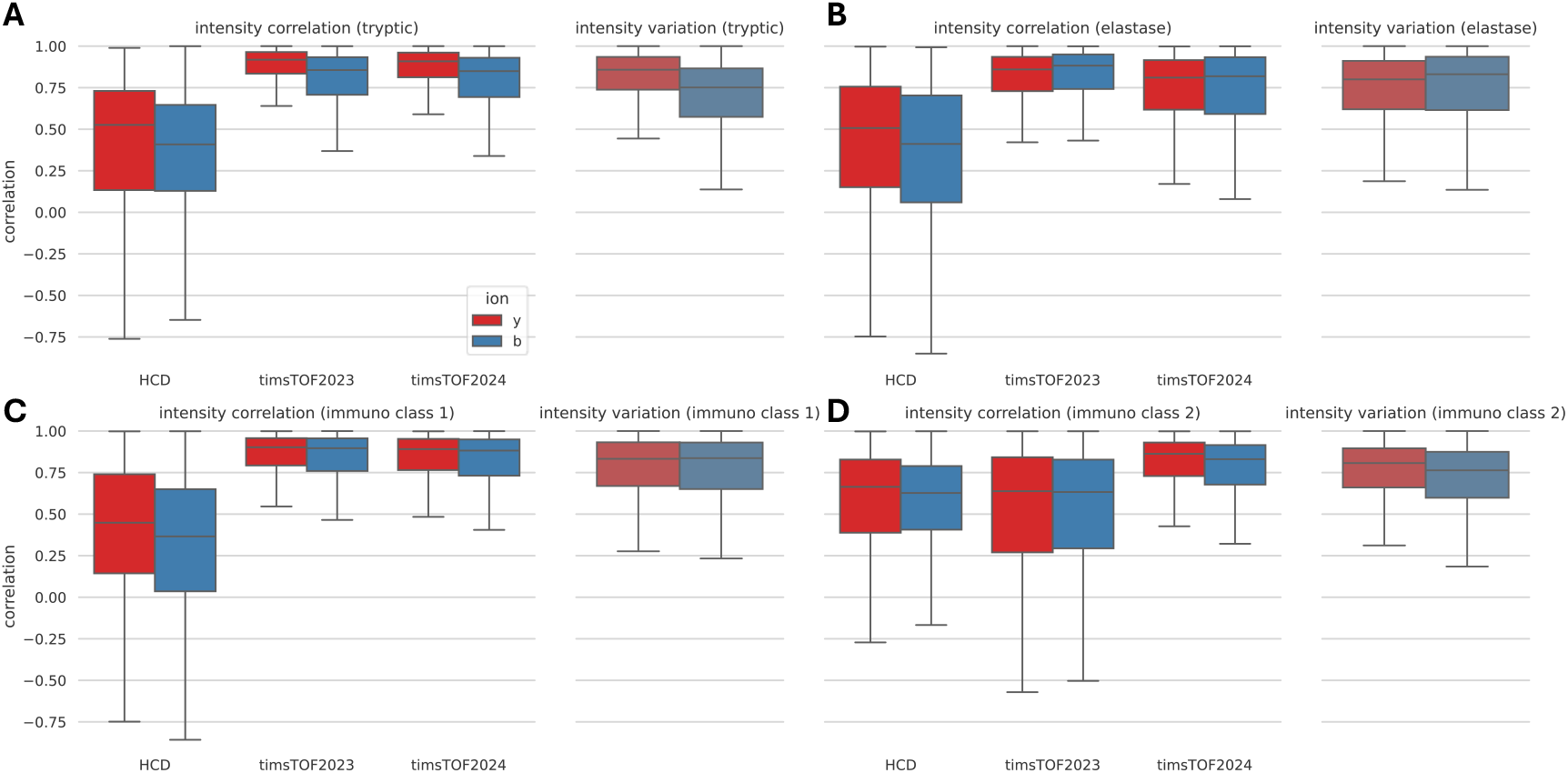
Boxplots showing intensity correlations with predicted and observed intensities for different prediction models. Also shown is correlation between observed intensities, roughly indicating maximal theoretical performance for the predictions. Four different peptide subgroups are analyzed: (a) tryptic peptides, (b) elastase digested peptides, (c) class I immunopeptides, and (d) class II immunopeptides.

## IM2Deep

### IM2Deep performance on modified peptides

To assess the CCS prediction ability of IM2Deep across a variety of differently modified peptides, we systematically evaluated its performance on all twenty-one modifications within the combined data set. Our approach involved training and optimizing twenty-one individual IM2Deep models, each exclusively trained on peptides not carrying one specific modification. These models were subsequently tested on peptides that do carry the excluded modification. Furthermore, we created two test scenarios: One where the excluded modification was encoded, and another where it was ignored. By comparing the prediction performance for both test scenarios, we gauged the ability of IM2Deep to predict the CCS for peptides with modifications unseen during training. This comparison aims to measure the improvement provided by IM2Deep over a basic approach that simply disregards the presence of the modification.

The prediction errors for each of the omitted modifications during training (Figure 2A) show an overall performance improvement when modifications are encoded during prediction. The observed reductions in mean absolute error (MAE) stem from IM2Deep’s accurate prediction of the CCS shift induced by the respective modifications, even though they were unseen during training. In addition to the reduced MAE, a general improvement in PCC is also observed. For example, in the case of formyl, an increase in PCC from 0.975 to 0.992 can be observed when encoding the modification in the test set (Figure 3, Supplementary Figure S2). This indicates that IM2Deep models not only learned the overall shift in CCS caused by modifications, but also captured how this shift depends on the specific context of the modification within each peptide.

**Figure 2:**
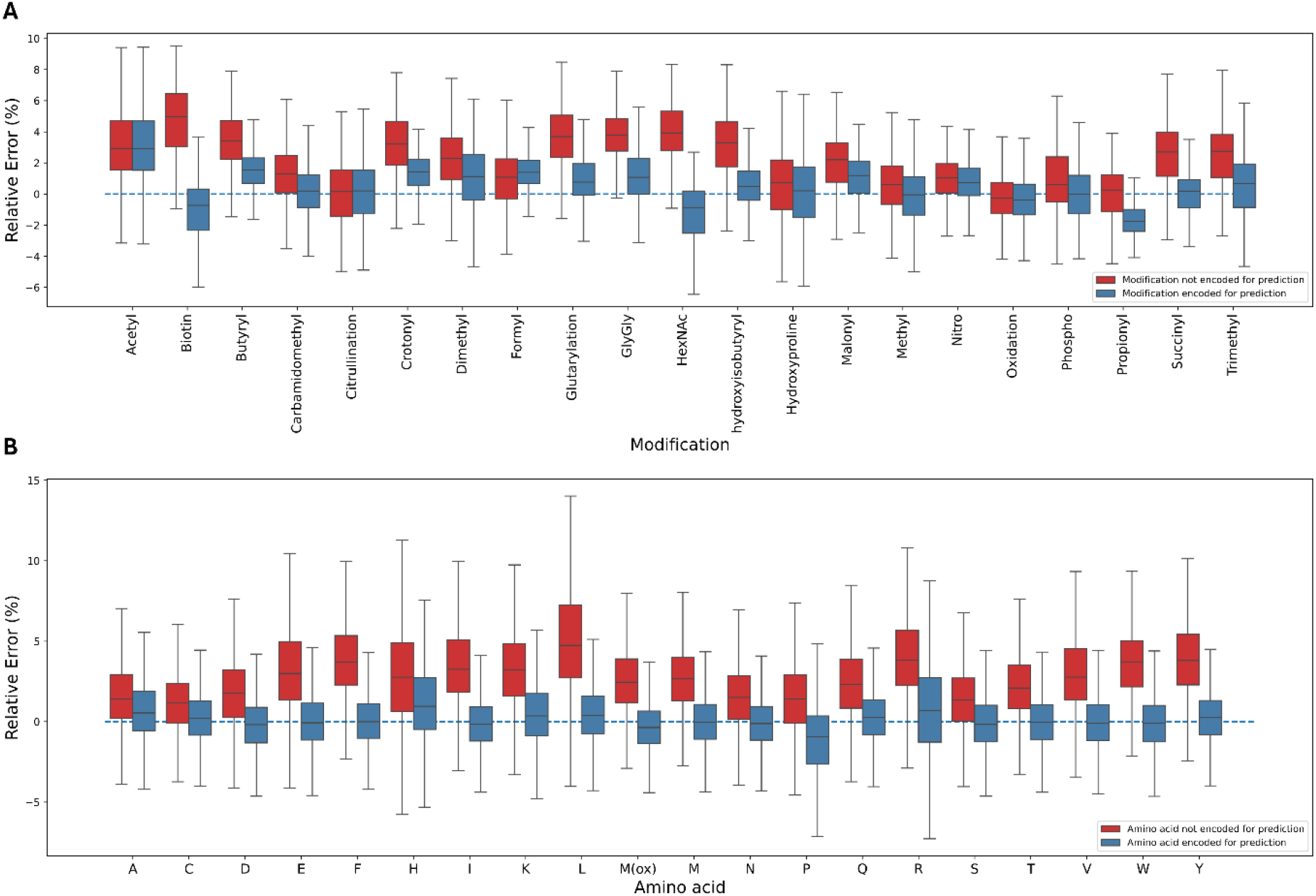
Our approach involved training individual IM2Deep models, each exclusively trained on peptides not carrying a specific modification/not containing a specific amino acid. The box plots show the IM2Deep prediction errors for peptides with modifications (A) and amino acids (B) that were not seen during training each of the respective models. Horizontal axis represents the excluded modifications/amino acids, while the vertical axis depicts the absolute error between the observed and predicted CCS when the modification/amino acid was either not encoded (red) or encoded during predictions (blue). These results indicate that IM2Deep generalizes well across modifications and amino acids, even if these were not seen during training. Note that peptides containing cysteine have a fixed carbamidomethyl modification.

**Figure 3:**
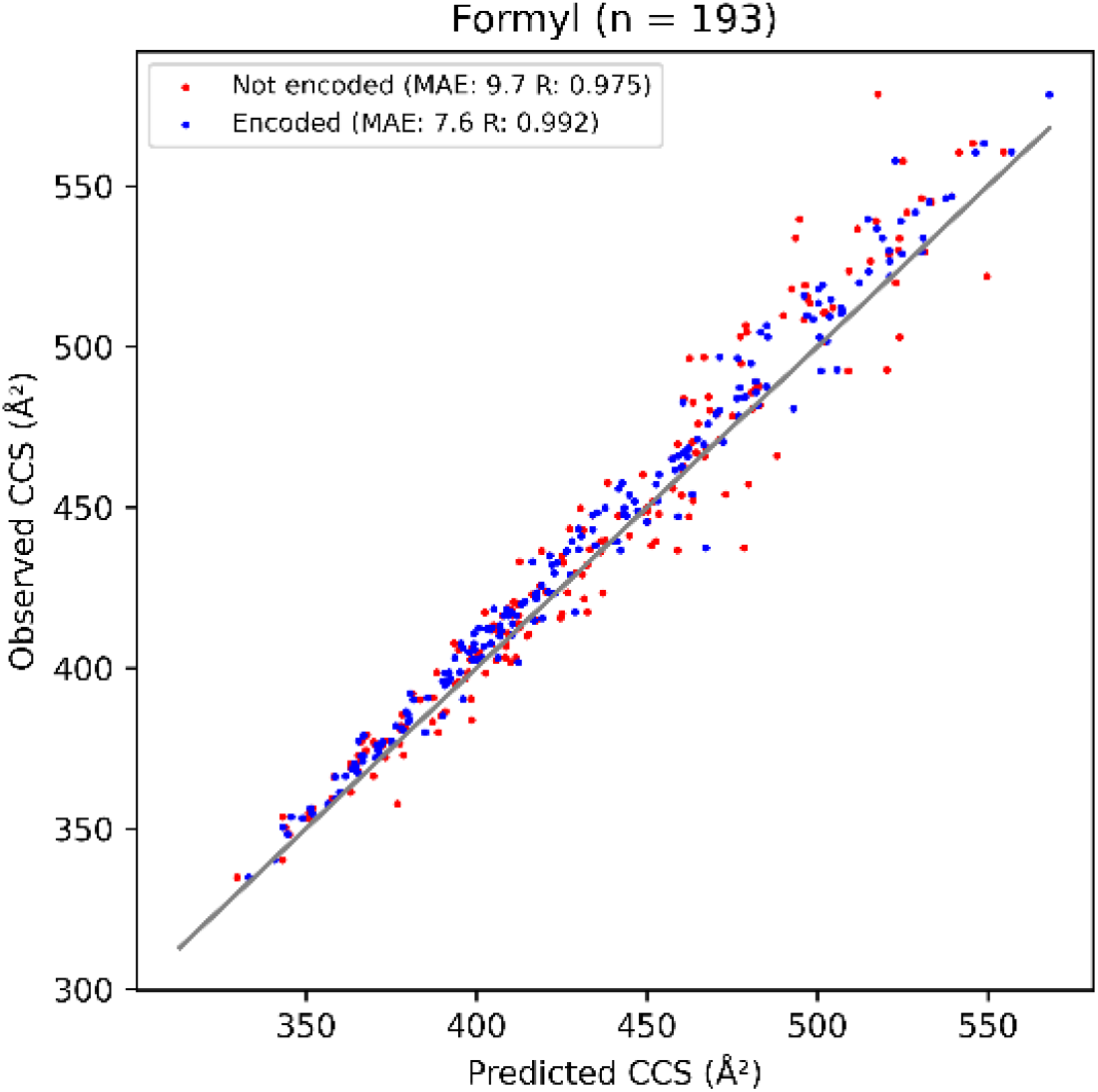
Scatterplot illustrating the performance of the IM2Deep model not trained on formylated peptides in predicting CCS for formylated peptides. The model was evaluated both with the modification encoded (blue) and ignored (red). Besides an improvement in MAE, an increase in Pearson correlation coefficient (R) is also seen. Variable n denotes the total number of formylated peptides.

In a second evaluation procedure, the same data set was used to train nineteen distinct IM2Deep models, each exclusively trained on peptides that lack a specific amino acid. Subsequently, each model underwent evaluation on peptides that did contain the amino acid excluded during training. Here too, two distinct test sets were generated from these remaining peptides: one where the excluded amino acid was encoded with its actual composition, and another where its composition was substituted with that of glycine. In this analysis, all peptides carrying modifications apart from methionine oxidation and cysteine carbamidomethylation were removed to focus on the performance on unseen amino acids. We demonstrate that encoding an amino acid as its own entity rather than as glycine lowers the MAE and increases the PCCfor most amino acids (Figure 2B, Supplementary Figure 3). Note that the diminished performance observed on proline is expected, as it can be attributed to its unique cyclic structure, which cannot be generalized from any of the other amino acids.

It is crucial to note that both of these evaluations are very stringent as the trained model has never encountered the respective modification or amino acid on which it is being evaluated. Furthermore, for the second evaluation, peptides that are similar and thus likely to contain the excluded amino acid will be collectively omitted from training, adding to the challenge. This factor is especially pertinent for lysine and arginine because all peptides in these data sets are tryptic. The resulting bias in training sets could potentially impact the model’s generalization ability. Nevertheless, despite these challenges, our model demonstrates high accuracy in predicting CCS values for amino acids absent from its training data, indicating its robustness and flexibility.

### Performance of the IM2Deep model shipped with TIMS^2^Rescore

A combination of the aforementioned evaluation data set and an immunopeptide data set (21) was used to train and evaluate the final IM2Deep model which is shipped with TIMS^2^Rescore. This data set was included to enhance IM2Deep’s performance on non-tryptic peptides, and on peptides with charge states 1, 5 and 6. Evaluation on the test set (10%) shows a mean absolute error of 6.26 Å^2^, a median relative error of 0.91% and a PCC of 0.996 (Supplementary Figure S4). Good predictive performance is observed across all charge states present in the dataset (Figure 4). The somewhat diminished performance on peptides with charge 6 can be explained by the limited number of training peptides (n = 249) for this charge state.

**Figure 4:**
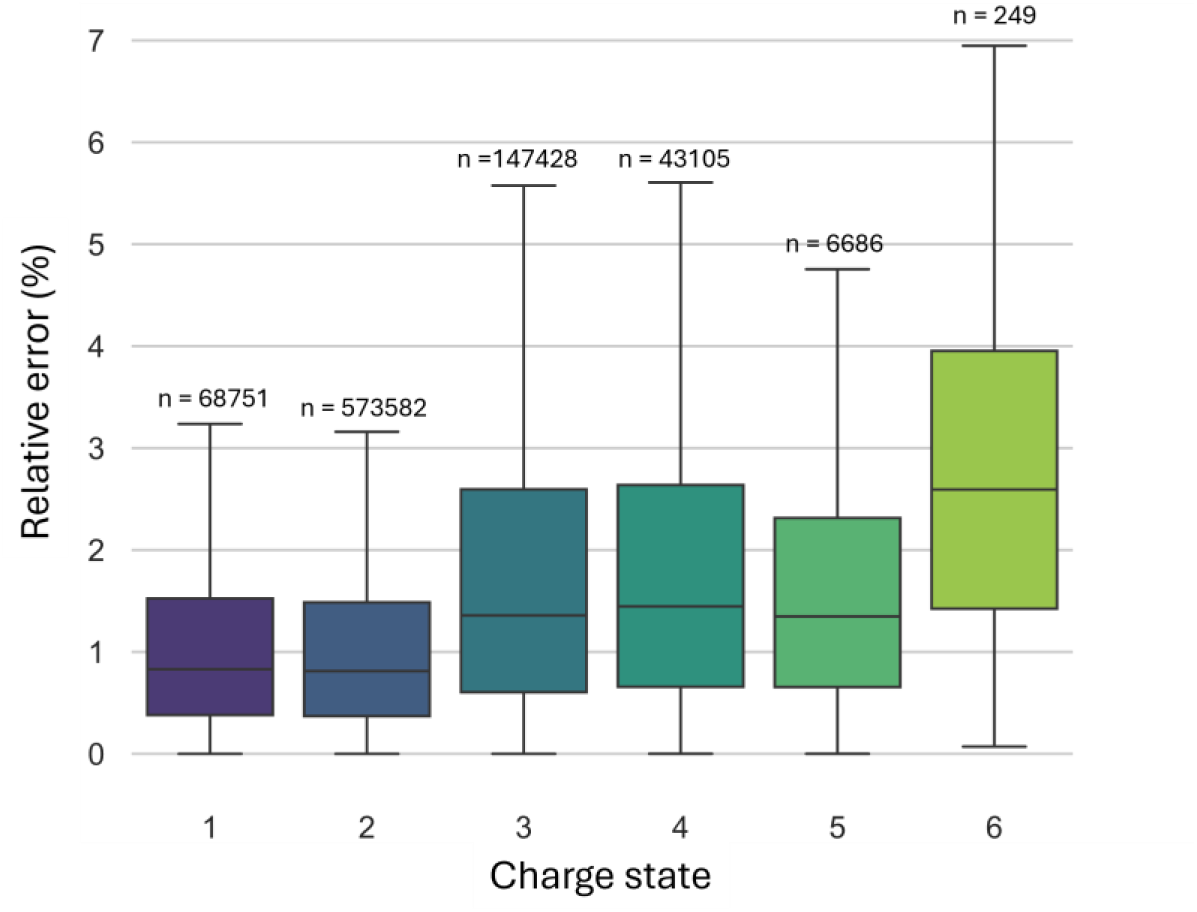
Box plots illustrating the relative error of the predictions made by the main IM2Deep model for peptides with different charge states. Numbers above the boxes indicate the number of training peptides with the corresponding charge state.

### Rescoring performance

Data coming from various proteomics subfields were used to assess (i) the overall rescoring performance of TIMS^2^Rescore, and (ii) evaluate the performance of the newly trained prediction models. Overall, we observe at least a 10% increase in confidently identified PSMs at 1% FDR and a 20% increase at the 0.1% FDR compared to Sage for all data sets (Figure 5). Most notably, we see a very large increase of almost 71% (191% at 0.1 % FDR) for the class I immunopeptides, which could be partially explained by the very large amount of data used for simultaneous rescoring, compared to the other data sets. Indeed, while Sage already identified 758,096 PSMs prior to rescoring for class I immunopeptides, this was only 83,412, 177,862 and 214,261 for plasma, meta proteomics, and class II immunopeptides respectively. Similar gains on the peptide level are also seen for all data types, where only the plasma proteomics data set has a slightly lower increase in peptide identifications than in PSMs. This is due to the few highly abundant proteins generating repeated spectrum identifications for the same peptides, such as albumin. Nevertheless, the newly identified peptides still lead to an 8% increase in plasma protein identifications. Similarly, for a highly complex sample such as gut metaproteomics, we observe a substantial increase in protein identifications. Moreover, the rescoring features from the new prediction models (PCC for the MS^2^PIP model and CCS error for IM2Deep), consistently show high correlations and low absolute errors, respectively, for confidently identified PSMs (Supplementary Figure S5-6). The former also results in higher feature weights in the rescoring algorithm, showing the importance of the new MS^2^PIP models. The latter, however, has smaller feature weights, despite the low errors, indicating that these features have a lower impact on rescoring. Most likely, this is due to the low orthogonality of CCS values with other features such as m/z and charge, as was previously described (29). Nevertheless, these features could still prove useful to further boost PSMs separation.

**Figure 5:**
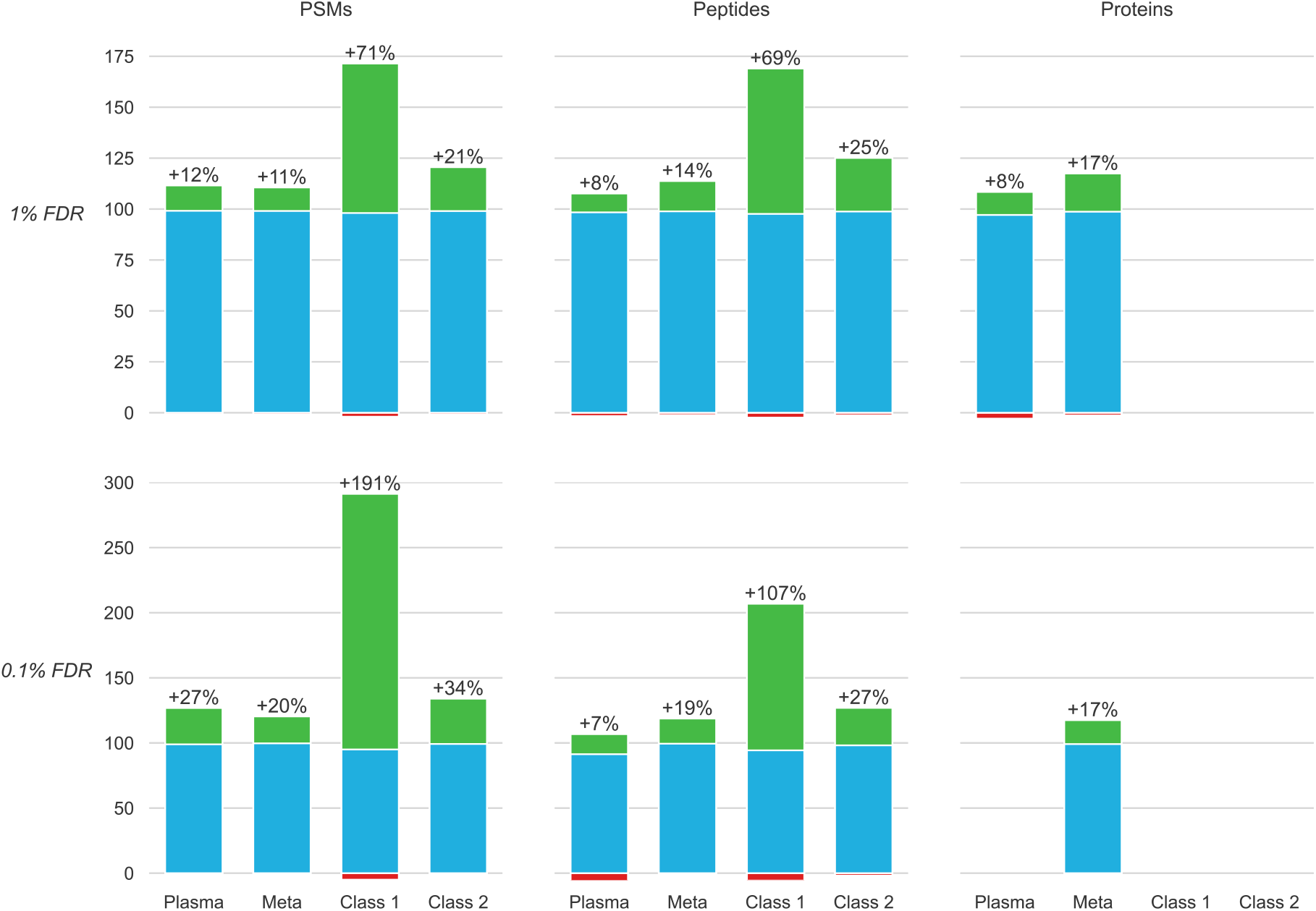
The gained (green), shared (blue) and lost (red) identifications at 1% and 0.1% PSM-, peptide-, and protein-level FDR, for Sage with TIMS^2^Rescore versus Sage without rescoring. Results for four data set types are shown: Plasma proteomics (Plasma), gut metaproteomics (Meta), class I immunopeptides (Class I), and class II immunopeptides (Class II).

## Discussion

Data-driven PSM rescoring is becoming a part of routine data analysis pipelines and has been repeatedly shown to greatly improve identification sensitivity and specificity (12–14) Simultaneously, state-of-the-art mass spectrometers, such as timsTOF instruments, provide increasingly higher sensitivities, pushing forward challenging proteomics subfields such as immunopeptidomics, plasma proteomics, and metaproteomics. Combining both technologies results in a much more performant analysis of a sample’s proteome.

We present a data-driven rescoring workflow optimized for timsTOF instruments. It includes new MS^2^PIP spectrum prediction models that accurately predict fragmentation behavior in timsTOF instruments for a wide range of peptide types. While the intensity correlations are generally lower than those expected for HCD Orbitrap spectra, they reach levels similar to the intensity variation observed between experimental timsTOF spectra. Moreover, these predictions do provide the expected boost in identifications when used in for PSM rescoring. This is reflected in the weights of the MS^2^PIP-derived features in the rescoring model (see Supplementary Figure S7). Furthermore, we included CCS predictions to further remove ambiguity for harder to identify PSMs, as previously shown for singly charged immunopeptides (29). With TIMS^2^Rescore, we provide a straightforward interface to the specialized timsTOF models for data-driven rescoring with direct support for Bruker raw spectrum files, thus alleviating the need for often slow and cumbersome file conversion steps. Together with the support for Sage (26), PEAKS (30) and ProteoScape PSM files, these new features make TIMS^2^Rescore an ideal post-processing tool for timsTOF identification pipelines.

We showcased the performance of TIMS^2^Rescore on several publicly available data sets of class I and class II immunopeptides, plasma proteomics, and metaproteomics experiments. Overall, TIMS^2^Rescore lead to gains of at least 10% in confidently identified PSMs, with more drastic gains for the larger and more challenging data sets, such as the class I immunopeptides. The plasma proteomics and metaproteomics data sets show similar gains at the protein level with 8% and 17%, respectively.

## Conclusion

Recent advancements in proteomics, such as highly sensitive mass spectrometers and the integration of AI, are clearly pushing the field forward. These technologies have made it possible to identify low abundant peptides and proteins with higher specificity, enhancing our understanding of biology and disease mechanisms. The development and application of tools like TIMS^2^Rescore with the newly trained MS^2^PIP and IM2Deep models demonstrate the positive impact of combining cutting-edge instrumentation with computational innovations. As we continue to leverage these advancements, the potential for new discoveries and improvements in disease diagnosis and treatment is vast.

## Availability

TIMS^2^Rescore is freely available and open source under the permissive Apache-2.0 license. It is available as a specialized command in the MS^2^Rescore Python package and is distributed through PyPI, Bioconda, and Biocontainers. The source code is available on GitHub at https://github.com/compomics/tims2rescore.

IM2Deep is open source under the permissive Apache-2.0 license and is freely available within TIMS^2^Rescore and as a stand-alone Python package on PyPI. The source code is available on GitHub at https://github.com/compomics/IM2Deep.

All data and scripts required to reproduce the presented results is available on Zenodo at https://doi.org/10.5281/zenodo.11277943.

## Author contributions

Arthur Declercq: Conceptualization, formal analysis, funding acquisition, investigation, methodology, software, validation, visualization, writing – original draft

Robbe Devreese: Conceptualization, formal analysis, funding acquisition, investigation, methodology, software, validation, visualization, writing – original draft

Jonas Scheid: Data curation, investigation, validation

Caroline Jachmann: Software

Tim Van Den Bossche: Data curation, investigation

Annica Preikschat: Data curation, investigation, writing – review & editing

David Gomez-Zepeda: Data curation, investigation, writing – review & editing

Jeewan Babu Rijal: Data curation, investigation, validation

Aurélie Hirschler: Data curation, investigation, validation

Jonathan R Krieger: Data curation, validation

Tharan Srikumar: Data curation, validation

George Rosenberger: Data curation, validation, writing – review & editing

Dennis Trede: Data curation, validation

Christine Carapito: Data acquisition, funding, supervision

Stefan Tenzer: Funding, supervision, writing – review & editing

Juliane Walz: Data acquisition, funding, writing – review & editing

Sven Degroeve: Conceptualization, methodology

Robbin Bouwmeester: Conceptualization, methodology, software, supervision, writing – review & editing

Lennart Martens: Conceptualization, funding, supervision, writing – review & editing

Ralf Gabriels: Conceptualization, methodology, software, supervision, writing – review & editing

## Supplementary Methods

### Immunopeptide class II data set generation

#### Cell culture and harvesting

The human B lymphoblastoid cell line JY (CVCL_0108) was purchased from ATCC. Cells were maintained in RPMI1640 medium supplemented with 10 % FCS (Gibco (v/v)), 1 mM sodium pyruvate, 100 units/ml penicillin, and 100 µg/ml streptomycin. Cells were harvested at 220 x g for 10 min, washed three times with 1x PBS prior to counting, and frozen at −80°C until further use.

#### Immuno-affinity purification of HLA peptide ligands

HLA class II ligands were enriched by immunoprecipitation as described by (31) with modifications (32). Briefly, the cell pellets were thawed and lysed in a non-denaturant buffer (1% CHAPS in PBS (m/v)) aided by sonication. Immunoprecipitation was performed using the anti-HLA-DR antibody L243 immobilized on CNBr-activated beads (Cytivia). The monoclonal antibody was purchased from Hoelzel-biotech and produced by Leinco Technologies (ref. H261). Samples were incubated overnight with the Antibody-beads, then washed once with PBS and once with water. Then, peptide ligands were eluted using 0.2% TFA (v/v) in water. Next, peptides were ultrafiltered using molecular weight cutoff (MWCO) filters (Vivacon 500, 10,000 MWCO Hydrosart, Sartorius). The flow-through was desalted by SPE on a Hydrophilic-Lipophilic-Balanced sorbent (Oasis HLB 96-well µElution Plate, 2 mg Sorbent per Well, 30 µm, Waters Corp.), applying 35% ACN (v/v), 0.1% TFA (v/v) for elution. The eluates were dried in a vacuum concentrator and dissolved in 15 µL of water with 0.1% FA (v/v) for subsequent LC-MS/MS analysis.

#### LC-MS analysis for HLA-DR immunopeptidomics profiling of JY cells

NanoLC-MS analysis was performed using a nanoElute coupled to a timsTOF-Pro-2 mass spectrometer. The desalted peptides were directly injected in a C18 Reversed-phase (RP) Aurora 25 cm analytical column (25 cm x 75 µm ID, 120 Å pore size, 1.7 µm particle size, IonOpticks, Australia) and separated using a 47 min gradient increasing the proportion of phase B (ACN with 0.1% FA (v/v)) to phase A (water with 0.1% FA (v/v)). The gradient started at 2% B, which increased to 17% within 23 min, then to 25% in the next 11.5 min, to 37% in 3.8 min, and to 95% in 3.8 min, before a wash step of 4.9 min at 95% B. A Captive Spray source was used for ionization, with a capillary voltage of 1600 V, dry gas at 3.0 L/min, dry temperature at 180 °C, and TIMS-in pressure of 2.7 mBar. Data was acquired using Compass Hystar and timsControl (Bruker) in DDA-PASEF mode using settings based on (32) . Ions were accumulated and resolved in 300 ms TIMS ramps from 0.65 to 1.75 Vs/cm^2^, using three MS2 frames per cycle and a cycle overlap of one. A stepped isolation polygon including singly charged ions above 445 m/z was designed to select precursors with a positive charge from 1 to 5 for fragmentation.. The fragmentation intensity threshold was set at 1000 and the target intensity at 20,000. The m/z acquisition range was set at 100 to 2000, and the high-sensitivity detection mode was activated.

## Supplementary Figures

**Figure S1:**
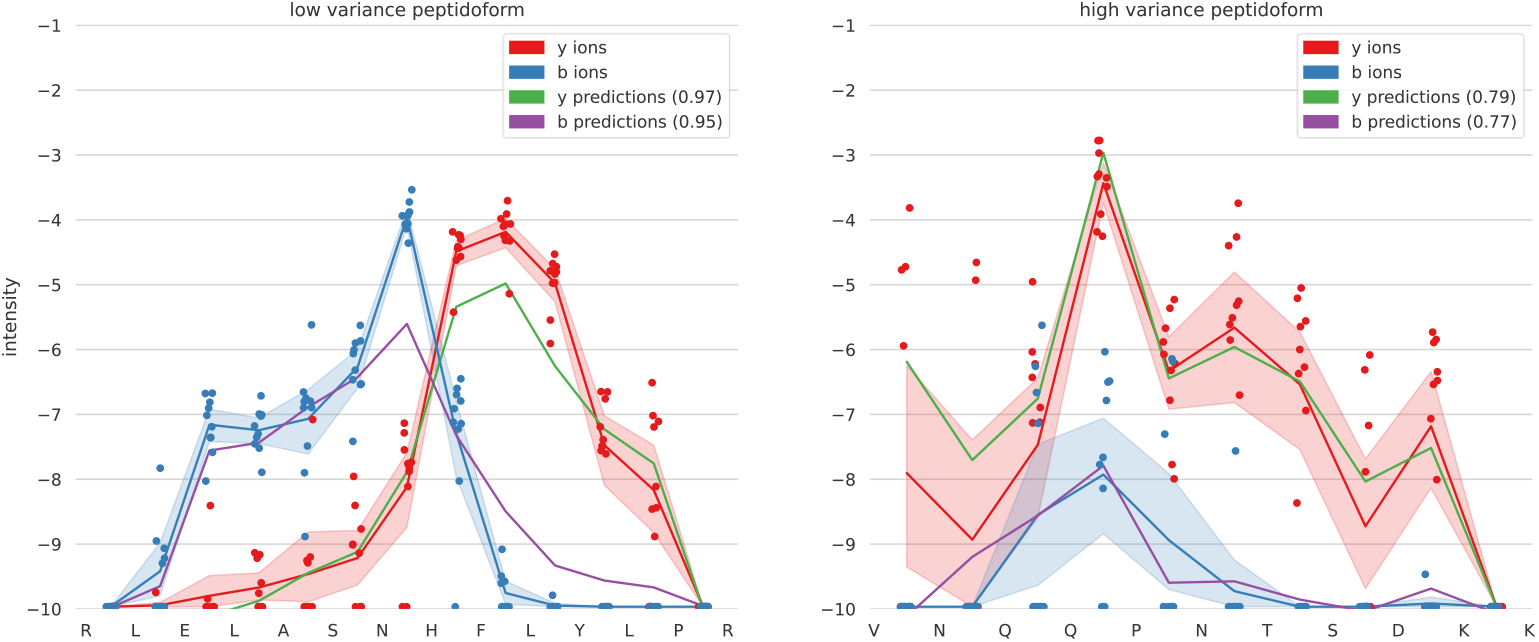
line plot showing the intensity variation for two different peptides, one with a low variation and one with a high variation for b (blue) and y ions (red). The predictions of the 2024 timsTOF model are depicted in green for the y ions and purple for the b ions.

**Figure S2:**
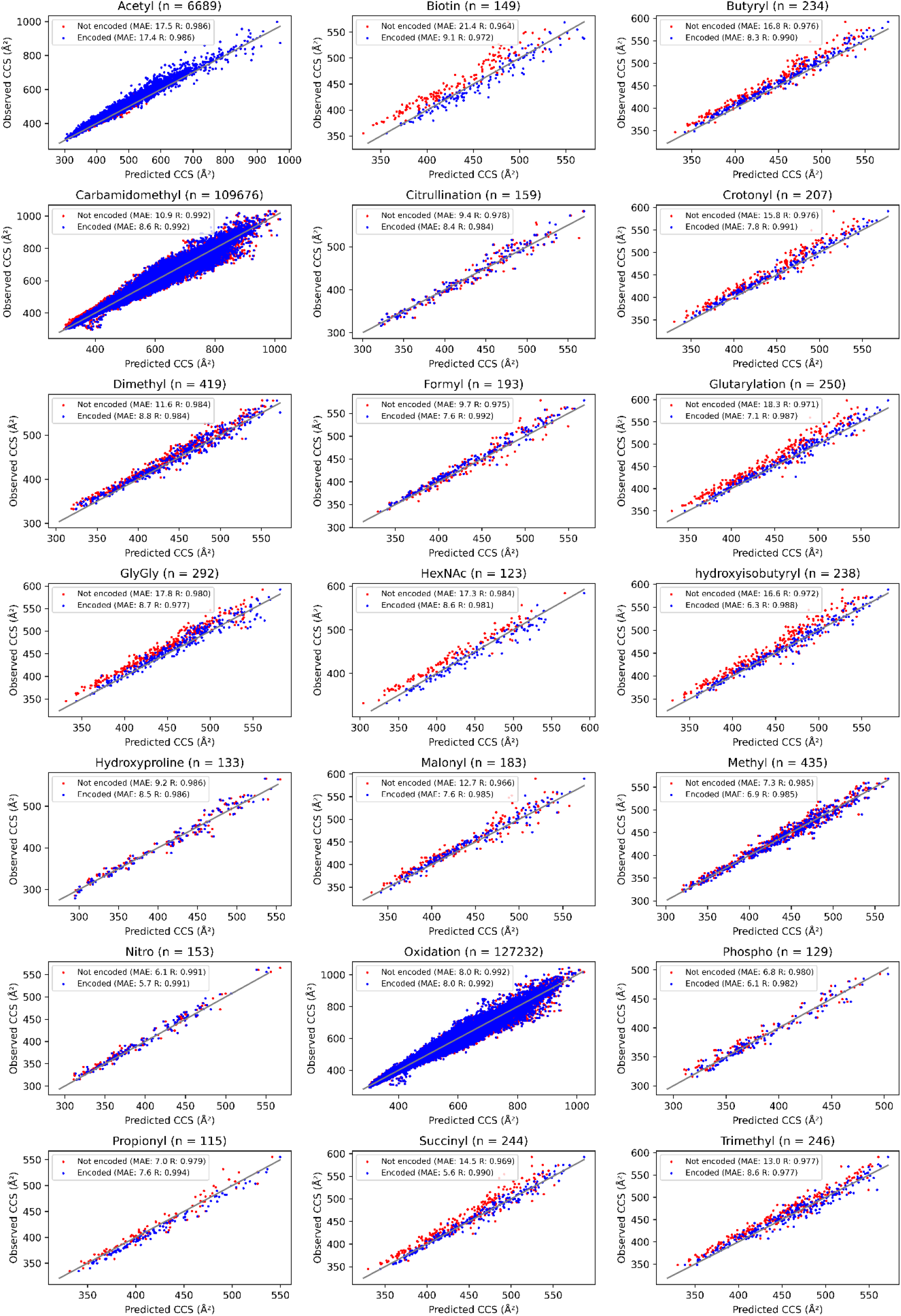
Each scatter plot displays the observed CCS plotted against the CCS predicted by models that were not trained on peptides carrying the specified modification. The dots represent CCS when modifications were either not encoded (red) or encoded (blue) by IM2Deep. N denotes the total number of peptides with the specified modification.

**Figure S3:**
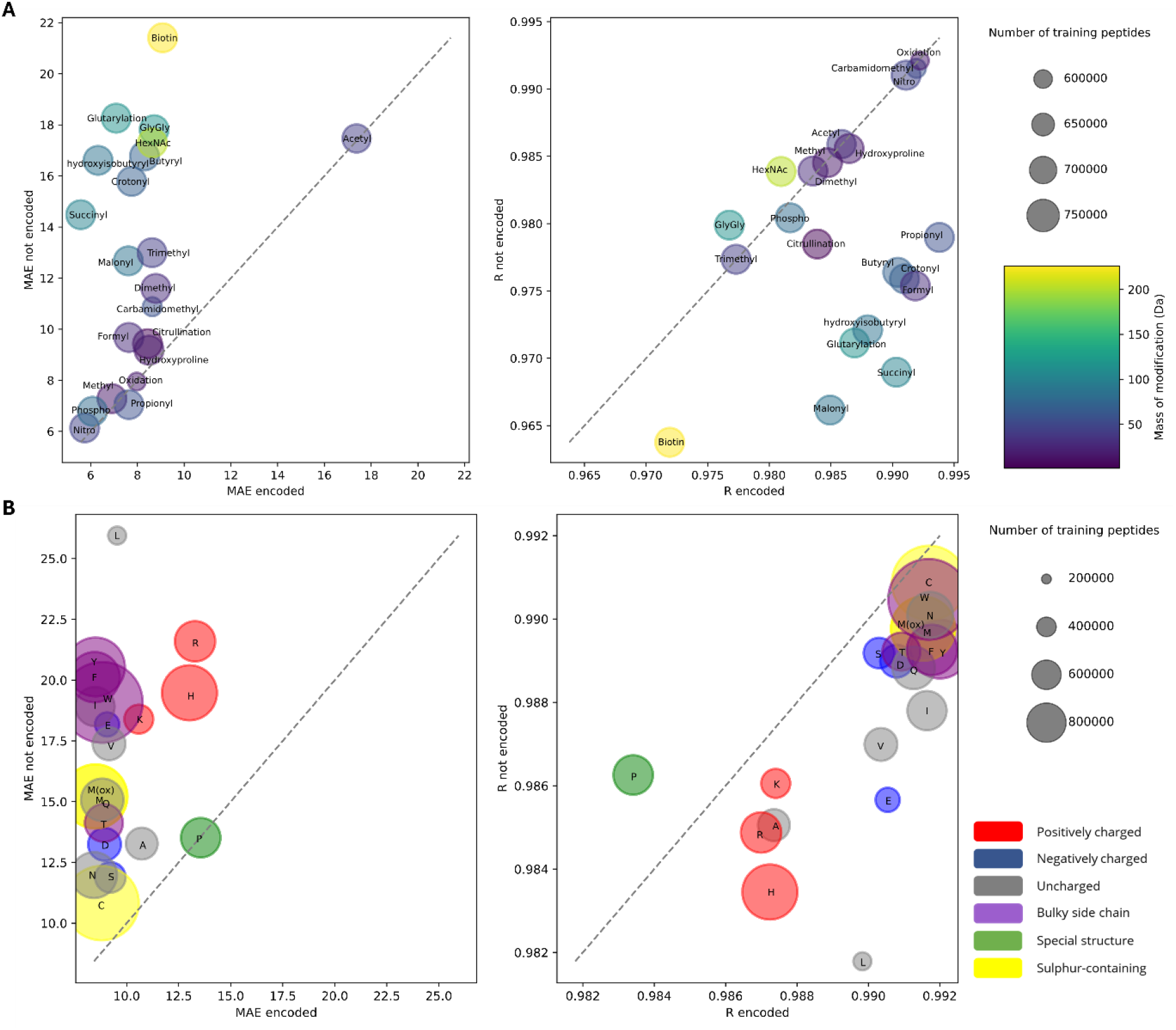
Each modification (A) and amino acid (B) excluded during training is represented as a circle, where the circle’s size indicates the remaining training peptides, and its color reflects the modification mass or the amino acid’s chemical characteristic. (A) The modification is either not encoded (vertical axis) or encoded with its atomic composition (horizontal axis). (B)The amino acid is encoded either as glycine (vertical axis) or as its own atomic composition (horizontal axis). Circle positions indicate the MAE (left) or Pearson R (right) for all modification-carrying or amino acid-containing peptides.

**Figure S4:**
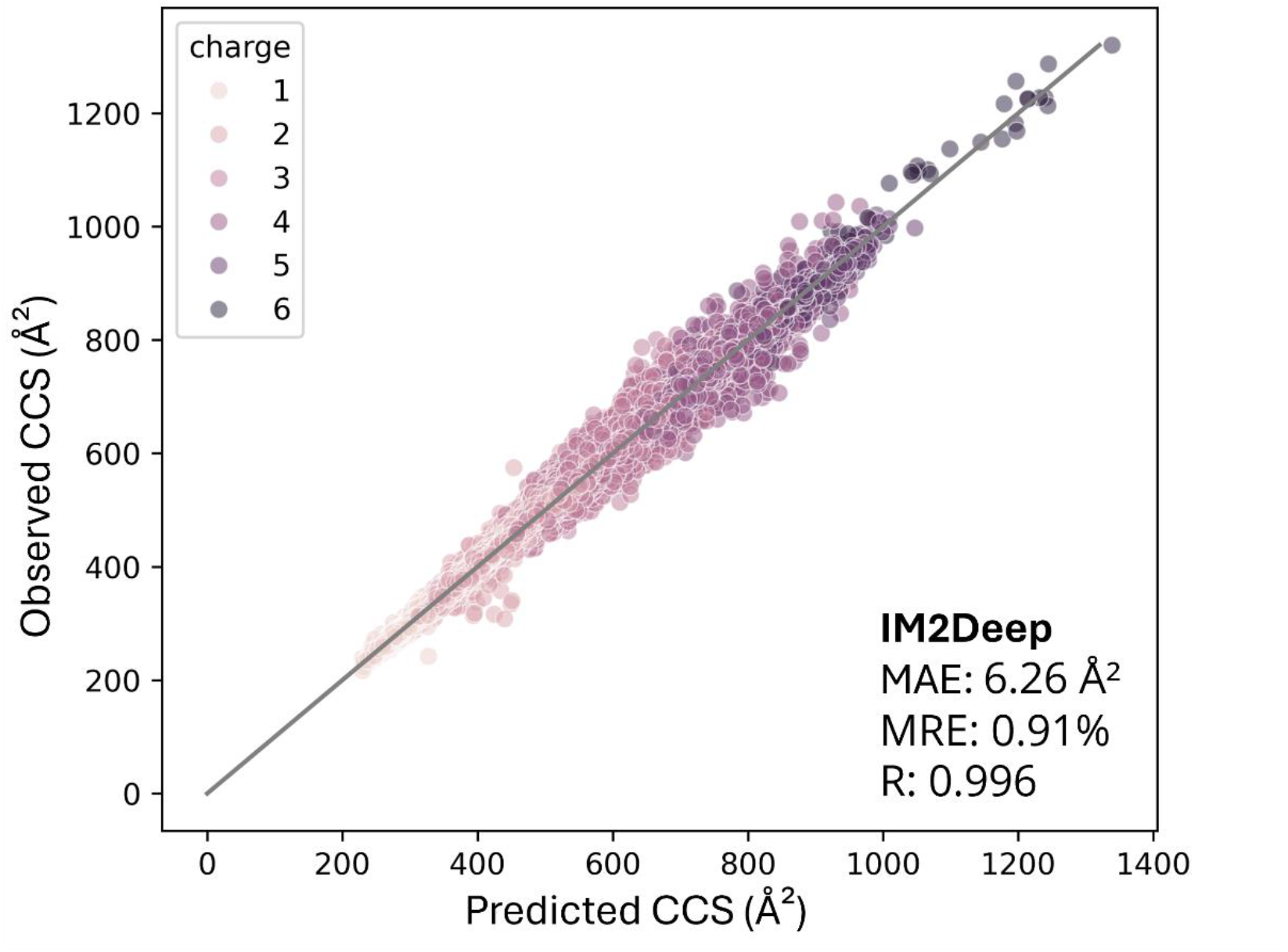
Test set performance of the base IM2Deep model, shipped with TIMS^2^Rescore. MAE: mean absolute error, MRE: median relative error, R: Pearson correlation coefficient.

**Figure S5:**
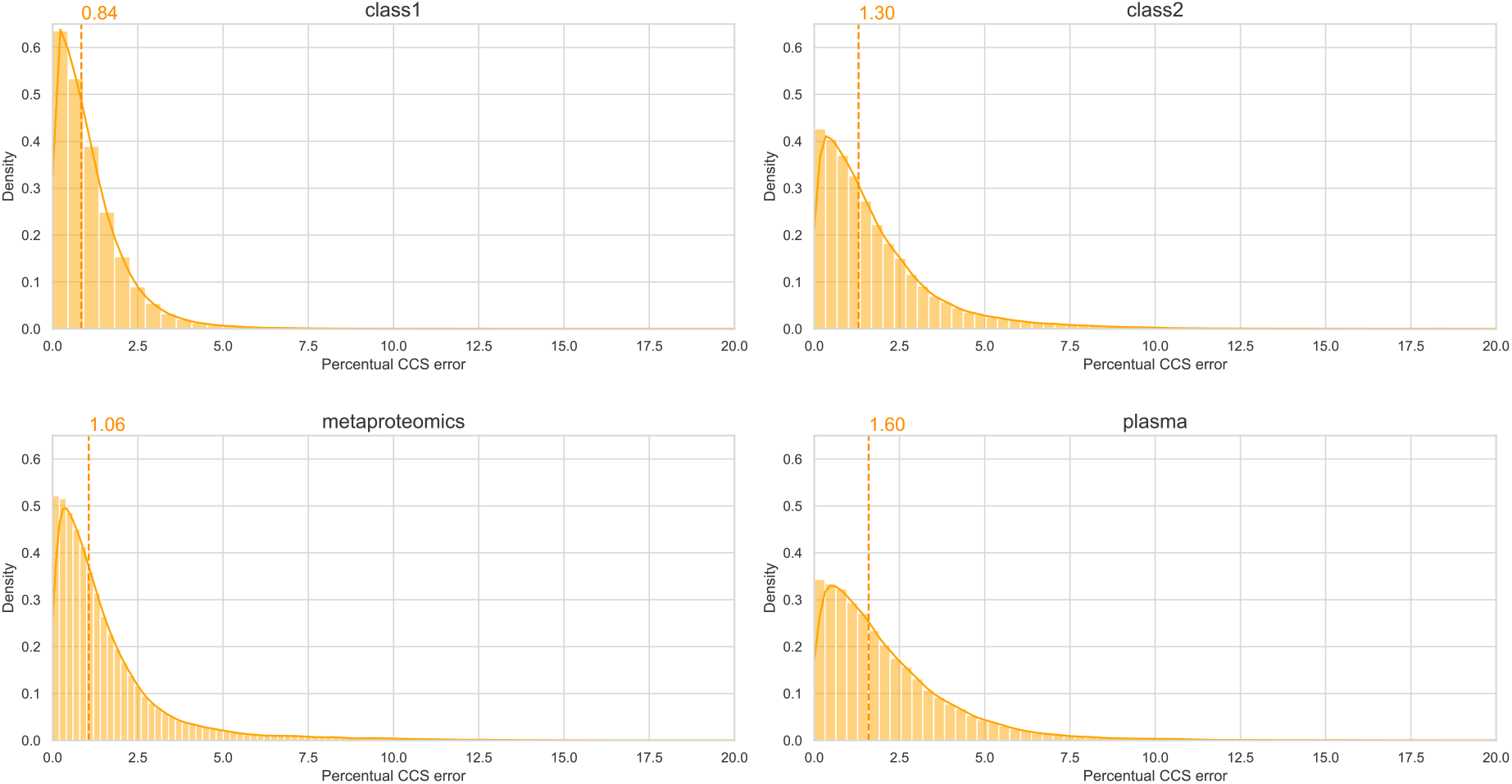
Kde plots showing the distribution of Percentual CCS error for all identified spectra after rescoring.

**Figure S6:**
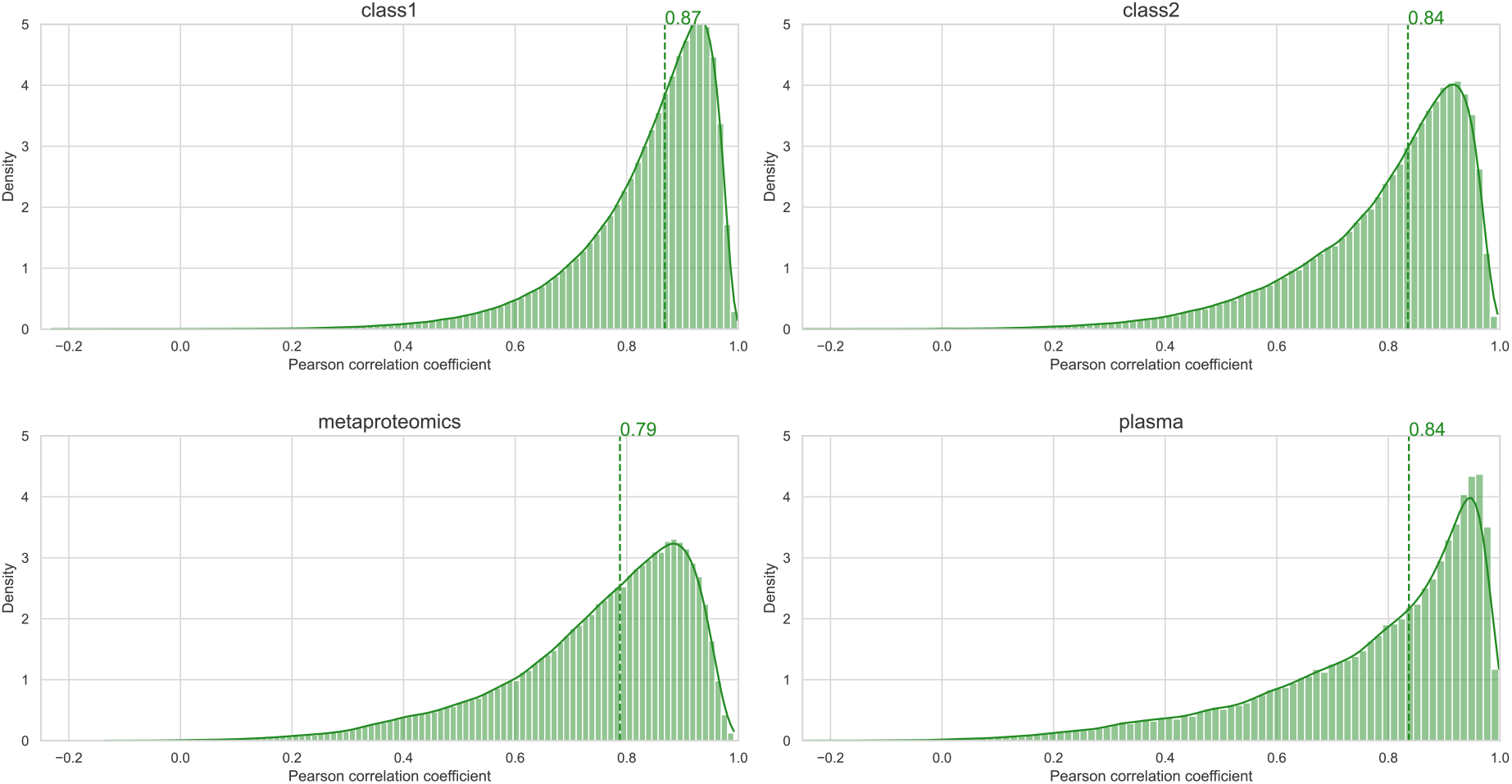
Distributions of Pearson correlation coefficients for all identified spectra after rescoring.

**Figure S7:**
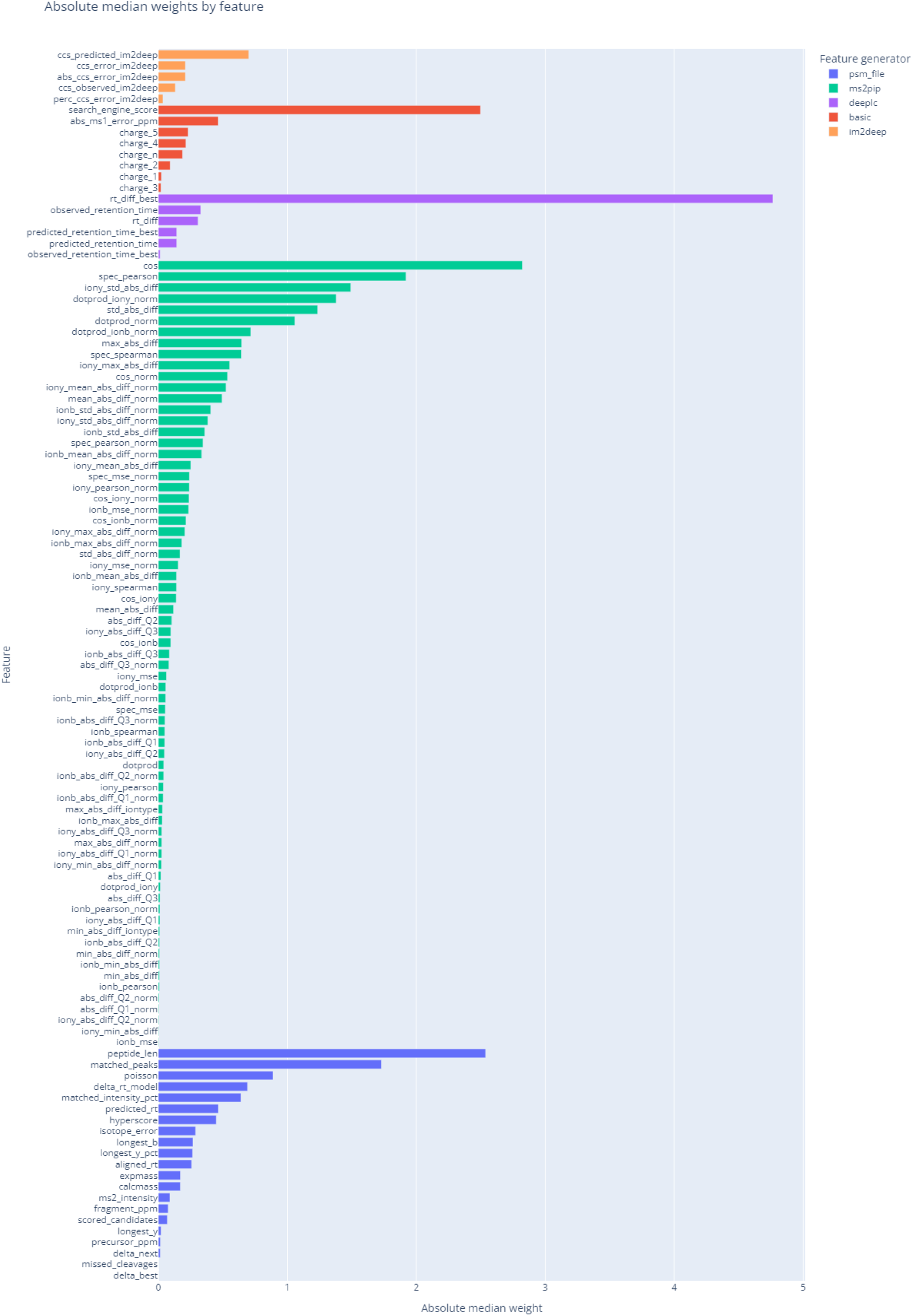
Feature weights for the class I immunopeptides rescoring run color-coded for different feature generators.

